# Hypoxia induced sex-difference in zebrafish brain proteome profile reveals the crucial role of H3K9me3 in recovery from acute hypoxia

**DOI:** 10.1101/2020.06.15.150052

**Authors:** Tapatee Das, Avijeet Kamle, Arvind Kumar, Sumana Chakravarty

## Abstract

Understanding the molecular basis of sex differences in neural response to acute hypoxic insult has profound implications for the effective prevention and treatment of ischemic stroke. Global hypoxic-ischemic induced neural damage has been studied recently under the well-controlled, non-invasive, reproducible conditions using zebrafish model. Our earlier report on sex difference in global acute hypoxia induced neural damage and recovery in zebrafish prompted us for comprehensive study on the mechanisms underlying the recovery. An omics approach for studying quantitative changes in brain proteome upon hypoxia insult following recovery was undertaken using iTRAQ-based LC-MS/MS approach. The results shed light on altered expression of many regulatory proteins in zebrafish brain upon acute hypoxia following recovery. The sex difference in differentially expressed proteins along with the proteins expressed in uniform direction in both the sexes was studied. Core expression analysis by Ingenuity Pathway analysis (IPA) showed a distinct sex difference in the disease function heatmap. Most of the upstream regulators obtained through IPA were validated at the transcriptional level. Translational upregulation of H3K9me3 in male led us to elucidate the mechanism of recovery by confirming transcriptional targets through ChIP-qPCR. The upregulation of H3K9me3 level in male at 4 hr post-hypoxia appears to affect the early neurogenic markers nestin, klf4 and sox2, which might explain the late recovery in male, compared to female. Acute hypoxia-induced sex-specific comparison of brain proteome led us to reveal many differentially expressed proteins, which can be further studied for the development of novel targets for better therapeutic strategy.

## INTRODUCTION

Oxygenation in vertebrates is always life or -death necessity for any of the metabolic needs of cells and tissues [1]. Over the last decade, we have acquired adequate information of cellular and molecular mechanisms in hypoxic -ischemic injury, survival and death [2, 3]. Hypoxic-ischemic neural injury continues to be the leading cause of death and disability worldwide [4]. The degree of disability does not simply reflect the severity or distribution of impaired blood supply [5]. The most common condition of hypoxia-ischemia leads to cerebral stroke due to the focal disruption of blood supply to a part of the brain. Other conditions include transient impairment of blood flow to the entire brain, termed global ischemia, which occurs following the cardiac arrest.

Low level of oxygen and the brain’s susceptibility to acute hypoxia depicts the key factor determining critical dependency. Cerebral oxygenation is reduced in hypoxia and neuronal damage can occur during a prolonged mismatch between oxygen supply and demand [6]. All the neurons in the brain can sense and, crucially modify its activity in response to hypoxia. Most neurons respond to hypoxia by decreasing metabolic demand and thus the need for aerobic energy [7]. Deciphering cellular response to energetic challenges that occur on the onset of acute hypoxia may give insight into the ischemic condition in various diseases [8]. Broad high throughput approaches in global changes in protein expression allows uncovering the critical signals underlying mechanisms in the disease condition. Acute Hypoxia causes a significant perturbation in cellular energy homeostasis before a hypoxia sensing and signal transduction cascade needing energy demand initiates [9]. An early component of the responses to acute hypoxia i.e. neural damage and recovery may have both post-transcriptional and translational mechanisms. The rapid response to acute hypoxia may preclude many pathways that require many new gene expressions suggesting the mechanism underlying recovery from acute hypoxia-mediated at least in part by the activities of the existing pool of mRNA and protein. An approach such as high throughput proteomic analysis is one of the ideally suited approaches to understand the neural changes induced by acute hypoxia with recovery [10].

Previous proteomic studies have shown hypoxia-induced changes in the zebrafish (*Danio rerio*) skeletal muscle proteome [11] and have implicated a broad range of cellular functions in response to hypoxia. Another proteomics study on zebrafish brain upon chronic unpredictable stress [12] has recently laid the groundwork for the analysis of neural proteome response to stressors.

A recent review article on Proteomics-Based Approaches for the Study of Ischemic Stroke [10] discussed the proteomics study of ischemic stroke using *in vivo* and *in vitro* models, with and without interventions and taking tissue, cerebrospinal fluid or plasma. Although proteomic studies have contributed with a long list of potential biomarkers for diagnosis, prognosis, and monitoring of ischemic stroke, most of these have not been implemented in clinical application successfully. The shortcomings from the existing proteomics data are small sample size, cell types, age of experimental animals, and using single-sex experimental animals seems to be responsible for blocking these results to achieve clinical implementation.

Like many neurological disorders, cerebral stroke too is reported to have sex-specific differences in occurrence and mechanisms. However, the molecular details underlying these sex-specific differences have not yet been explored using the relevant animal model. In fact, many factors including genetics, hormones (estrogen and androgen), epigenetic regulation, and environment contribute to sex-specific differences. Since ischemic sensitivity varies over the lifespan, and “ischemia resistant” female phenotype diminishes after the menopause, hence the role of sex hormones cannot be ruled out. To understand the role of hormonal status on the cerebral vasculature in pin-pointing sex-specific differences in stroke pathophysiology, a suitable animal model that can be simpler and helpful to address these complicated sex-specific differences is warranted.

Sex-specific differences in the hypoxic-ischemic brain have profound implications for effective prevention and treatment. Global hypoxic-ischemic damages and recovery are well studied under the well-controlled, non-invasive, reproducible conditions in zebrafish [13–17]. In our previous study, we have reported the sex-specific difference in hypoxia-induced neural damage and recovery, where we have concluded that as compared to male, female showed higher level of neural damage and an ability to recover faster. This interesting finding led us to explore the global proteome changes induced in recovery after the hypoxic stress, so in the present study, we performed a high throughput proteomic analysis on zebrafish brain by iTRAQ method. iTRAQ labelling method also allows to identify different post-translational modifications which are key to understand the aetiology and better treatment.

### Experimental Procedure

#### Protein extraction for iTRAQ

The animals were euthanatized and decapitated to remove the brain. The whole brain from each animal was homogenized in lysis buffer (50 mM ammonium bicarbonate pH 8.0, 0.1% SDS with protease inhibitor cocktail (Sigma) and for further efficient disruption and homogenization of tissue, a mild sonication was done using Bioruptor^®^. The obtained lysates were cold-centrifuged at 14,000 rpm for 15 min and the supernatant was quantified using Bradford assay with BSA as standard. Further protein samples were cleaned up by acetone precipitation. For each group 80 μg of protein was taken and six-volumes of chilled acetone were added for precipitation. After decantation of acetone the samples were air dried.

Before trypsin digestion, all the protein samples were reduced and cysteine blocked using the reagents provided in the iTRAQ^®^ Reagents-4plex Applications kit-Protein (AB Sciex). Digestion and labelling of proteins were done according to the manufacturer’s protocol. The samples from normoxia male and female were labelled with reagents 114 and 116 and the samples from hypoxia male and female were labelled with reagents 115 and 117, respectively. Subsequently, all the labelled samples were pooled and vacuum dried and further cleaned up using the C18 desalting column (Thermo Fisher Scientific). The final fraction was concentrated using a vacuum concentrator and reconstituted in 10 μl of 0.1% formic acid for LC-MS/MS analysis.

#### LC-MS/MS Analysis

LC-MS/MS analysis of the trypsin digested iTRAQ labelled and purified fractions were performed in LTQ - Orbitrap Velos (Thermo Scientific, Germany). The fragmentation was carried out using higher-energy collision dissociation (HCD) with 50% normalized collision energy. The MS data were analysed using Proteome Discoverer (Thermo Fisher Scientific, Version 1.4). MS/MS search was carried out using the SEQUEST search engine against the NCBI zebrafish protein database. Search parameters included trypsin as an enzyme with maximum 2 missed cleavage allowed; precursor and fragment mass tolerance were set to 10 ppm and 0.2 Da respectively; Methionine oxidation was set as a dynamic modification while methylthio modification at cysteine and iTRAQ modification at N-terminus of the peptide were set as static modifications. The FDR was calculated by enabling the peptide sequence analysis using a decoy database. High confidence peptide identifications were obtained by setting a target FDR threshold of 1% at the peptide level. Relative quantitation of proteins was determined based on the ratios of relative intensities of the reporter ions from hypoxia treated and untreated samples released during MS/MS fragmentation of each peptide. Appropriate quality control filters at the level of peptides/peptide spectral matches (PSMs) and then at the protein level were applied to the iTRAQ data. Proteins identified from the triplicate runs having more than 1.5-logfold changes in the hypoxia samples against the normoxia samples were selected for upregulation and having less than 0.5-log fold change considered to be down-regulated for its differential expression. Proteins based on their regulation were analysed for putative associations in different network pathways.

#### Protein enrichment analysis

To perform the functional enrichment tests of the candidate proteins, we used Ingenuity Pathway Analysis (IPA) software for both canonical pathways and molecular networks altered. The IPA system provides a more comprehensive pathway resource based on manual collection and curation. The rich information returned by IPA is also suitable for pathway crosstalk analysis as it has more molecules and their connections included. For analysis we have provided the identified peptides with relative and absolute expression fold change values and performed core IPA analysis, biomarkers, and molecular and functional comparison analysis.

#### Co-Immunoprecipitation

Zebrafish brain tissue was homogenized in nuclear extraction buffer (50 mM HEPES (pH 7.8), 50 mM KCl, 300 mM NaCl, 0.1 M EDTA, 1 mM DTT, 10% (v/v) Glycerol and 1X protease inhibitor) and further washed with PBS and incubated in RIPA buffer ≥20□mM Tris (pH□7.5), 150□mM NaCl, 1% NP-40, 5□mM EDTA, protease and phosphatase inhibitors) for 15□min on ice. After centrifugation, the supernatant was collected and pre-cleared with protein A agarose beads (Santa Cruz) at 4□°C for 30□min. The pre-cleared lysate was then incubated with Anti-Histone H3 (tri methyl K9) antibody (H3K9me3) (AB8898 1:250) complexed to protein A beads at 4□°C for 5-6□h, followed by washes with a buffer containing 10□mM Tris (pH□7.5), 150 mM NaCl, and 1 mM EDTA. The beads complexed with the immunoprecipitated proteins were then boiled at 100□°C in 3X Laemmli buffer for 5□min. 2.5% of whole tissue lysate was taken as input for each immunoprecipitation. Western blotting was carried out by loading equal amounts of the immunoprecipitated proteins.

#### Immunoblotting analysis

For immunoblotting experiments, cells were lysed in 3X Laemmli buffer (180□mM Tris (pH□6.8), 6% SDS, 15% glycerol, 7.5% β-mercaptoethanol, and 0.01% bromophenol blue). Images were captured using Chemicapt (Vilber-Lourmat, Germany). Densitometry analysis for blots was performed using ImageJ software (NIH) and images were processed in Adobe Photoshop CS3. The intensity values plotted or mentioned are average values from the number of biological replicates indicated in the legend.

#### Chromatin immunoprecipitation (ChIP) assay

ChIP was performed as described in [18] with required minor modifications. Briefly, for each ChIP, cross-linked samples from three animals were pooled together both in normoxia and hypoxia group. The 30 μg of chromatin from each sample was pre-cleared with Dynabeads (Invitrogen) before incubation with an *anti-rabbit* H3K9me3 antibody (EPR16601) keeping non-immune rabbit IgG antibody as a negative control. After reverse cross-linking, sequential washes with different concentration salt buffers, DNA was purified using phenol-chloroform-isoamyl alcohol (25:24:1 ratio, SIGMA). Specific primers for the gene-specific 5’ upstream region of the transcription start site were used for quantifying the enrichment of the histone mark H3K9me3, for 10 % input, in SYBR Green-based qPCR assays.

#### qPCR

The total RNA was isolated using TRIzol Reagent as per the manufacturer’s instruction. The cDNA was synthesized employing RevertAid H Minus First Strand cDNA Synthesis Kit according to manufacturer’s protocol. The primer sequences are available on a request basis. Real-time PCR was performed in triplicate using SYBR Green PCR Master Mix Detection System (Applied Biosystems). Normalization of mRNA expression levels was carried out using β-actin as the housekeeping gene.

## RESULTS AND DISCUSSION

### Analysis of zebrafish brain proteome induced by acute hypoxia and during recovery using iTRAQ based LC-MS/MS

As described previously by us [16], 4 hours post-acute hypoxia treatment (for 5 min at DO = ± 0.6 mg/liter) the brain tissues from both hypoxia-treated and untreated male and female zebrafish were isolated (n=6 per group) and subjected to comprehensive proteomic profiling. For this, male and female zebrafish brains of nine individuals from each gender were pooled together for iTRAQ based LC-MS/MS method and three technical replicates were run in LC-MS/MS, as depicted in Fig. 1A. Differentially expressed proteins were identified using iTRAQ and LC-MS/MS analysis on LTQ Orbitrap Velos mass spectrometer, by comparing hypoxia male (HM) vs normoxia male (NM) and hypoxia female (HF) vs normoxia female (NF) (Fig 1A). 2,323 proteins were identified to be regulated differentially after the data analysis from experimental runs in triplicate. Majority (i.e. 84%) of differentially regulated proteins, 1968 in total, were found downregulated in HM versus NM. In contrast, more than half (51%) of differentially regulated proteins, 1188 in total, were found upregulated in HF versus NF (Fig 1B). 1535 proteins were differentially regulated among those 1518 (98%) proteins were upregulated in female and downregulated in male whereas only 17 (2%) proteins were upregulated in male and downregulated in female showing a clear differential regulation in a sex-specific manner. 554 proteins showed similar expression pattern in both sexes where 478 proteins were upregulated and 76 proteins downregulated.

**Fig 1.**
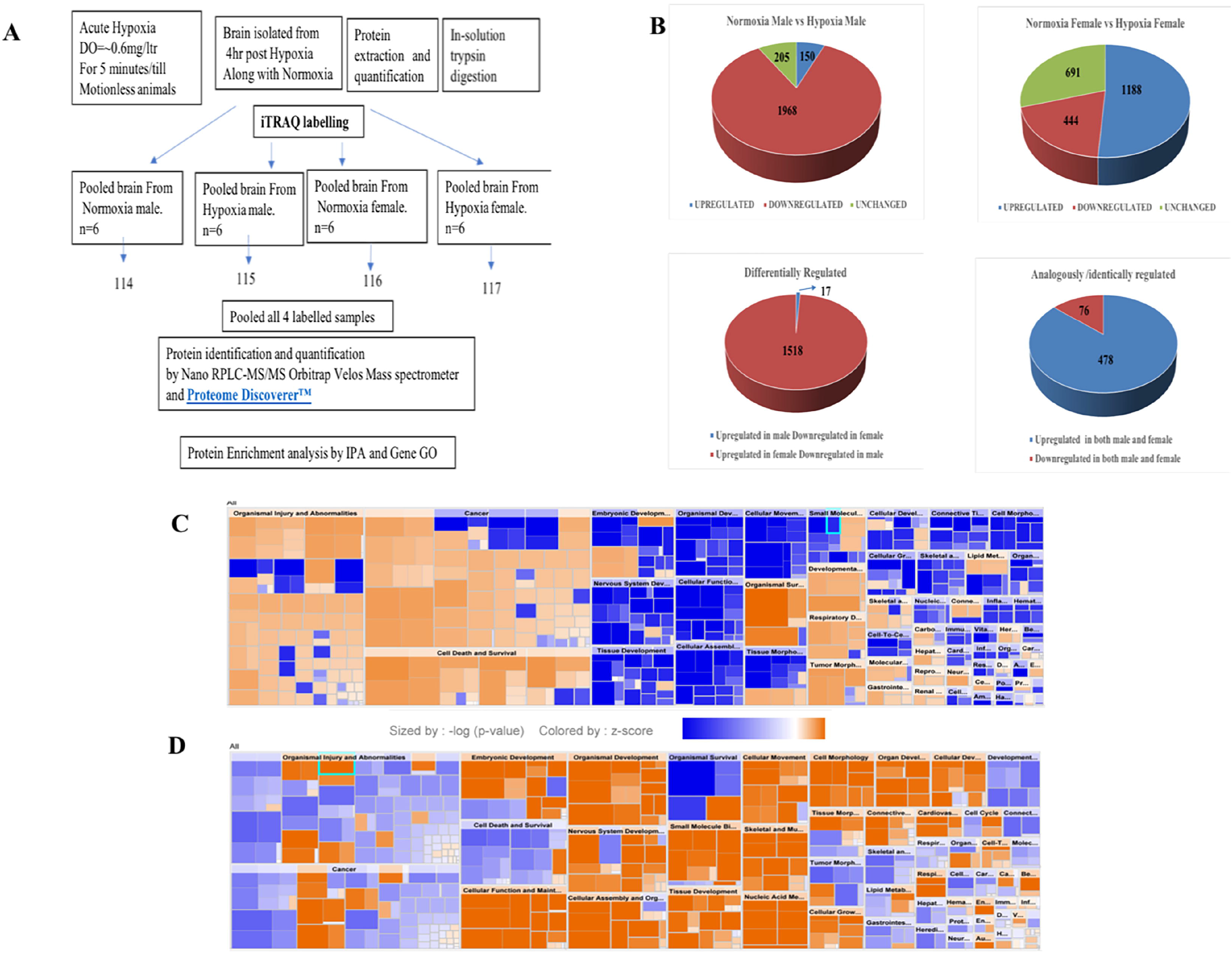
Analysis of zebrafish brain proteome by iTRAQ and Comparative heat map generated by IPA. Schematic representation of brain proteome analysis by iTRAQ labelling (A), Pie chart showing the analysis of expression of proteins resulted from iTRAQ in hypoxia male and female brain compared with normoxia male and female brain (B).Heat map of male (C) and female (D) zebrafish brain proteome upon hypoxia treatment.

### Protein enrichment analysis for zebrafish brain proteome induced by acute hypoxia and during recovery

The protein enrichment analysis on the altered proteome was performed using the Ingenuity Pathway Analysis (IPA) software. The disease function pathway-based heat map generated by the IPA (Fig 1 C-D) clearly showed a sex-specific difference in the altered expression of proteins in different disease pathway conditions.

The IPA analysis for disease function annotation showed a predicted activation state with activation z-score and p-values and molecules involved in each category of disease function. Among all the 502 proteins mapped in IPA for the disease and function analysis, in male 65 categories of disease function showed the predicted activation state: 37 categories showed increased activation state while 28 categories exhibited decreased activation state. In female only 30 categories of disease function showed the predicted activation state and among those 21 categories of disease function showed increased activation state and only 9 categories showed decreased activation state.

The most striking feature was the contrasting regulation in male and female in one of the disease and function categories named organismal survival and leads to organismal death with a very significant (p= 2.31E-11) activation z-score (12.434) identifying 251 molecules with increased activation state; but the same 251 molecules showed a significantly (2.59E-11) decreased activation state with a z-score (−7.796) in female (Table 1). In male most of the increased activation state was observed in organismal injury, abnormalities, cell death, connective tissue disorders, developmental disorder, skeletal and muscular disorders, neurological, respiratory diseases and cancer; whereas in female cellular assembly and organization, cellular function and maintenance, cell morphology, nucleic acid metabolism, cellular movement showed an increased activation state. The disease function analysis clearly showed a sex-specific difference in hypoxia-induced neural damage and recovery as seen in our earlier studies [16].

Based on the zebrafish annotated database IPA mapped 994 proteins out of 2323 proteins identified in the iTRAQ analysis. These 994 proteins included different types of proteins i.e. transporters, transmembrane receptors, translation and transcription regulators, phosphatases, peptidase, kinases, enzymes, G-protein coupled receptors, ligand-binding receptors, and cytokines (Table 4.2). Out of all the groups, the majority of proteins belong to the group “transcription regulators”. The upstream regulators analysed in IPA were 155 in male and 165 in female; among these 18 upstream regulators in male and 24 in female showed the predicted activation state (Table 4.3). Among the 18 upstream regulators in male 14 were inhibited and only 4 were activated and most of these were transcription regulators. Among the 24 upstream regulators in female as many as 16 were activated with majority of transcription regulators and just 8 were inhibited, which were not found in male. Five upstream regulators (Myc, Mknk1, Nfe2l2 (Nrf2), Thrb and Otx 2) were found to be common in both male and female with differential activation state, and interestingly these were in opposite direction i.e. inhibited in male while activated in female.

In Table 4.4 the regulator effects of male and female are shown, where only one of the regulators Nfe2l2 is common in both but with only one common target molecule VCP (valosin containing protein) and all different target molecules in both the dataset VCP was earlier reported to be an AKT binding protein and its expression was found enhanced in hypoxia [19].

Few of the target molecules (Eno1, Foxo1, Gp1, Hmox1, Nos2, Pkm, Ran, and Vcp) from both the data set were considered for validation by quantitative Realtime PCR (Fig 2). *eno1*(*enolase* 1) is one of the HIF target gene [20]. The qPCR analysis revealed more than ~2-fold increase in *eno1* and *ran* (ras-related nuclear protein) in female but remained unchanged in male. The foxo transcriptional factors are important regulators of cell survival in response to various stresses including oxidative stress [21]. *foxo1* was upregulated ~4-fold in female but unaffected in male thus indicating better survival response after hypoxia in female. The expression of *gp1* (Glycoprotein 1) and *hmox1* (Heme Oxygenase 1) showed similar kind of expression pattern in both sexes, a mild upregulation in male and ~3-fold upregulation in female. (*gp1*) act as a glycolytic enzyme besides it functions as a tumor-secreted cytokine and an angiogenic factor (AMF) that stimulates endothelial cell motility GPI is also a neurotrophic factor (Neuroleukin) for spinal and sensory neurons. Role of neurotrophic factor in repair mechanism are well evident, therefore in our study 4hr post-hypoxia female is better in recovery speed as compared to male. *Hmox1* hhas been shown to be induced by various stresses including hypoxia [22]; our study also revealed an increase in its expression. The expression of *nos2* (nitric oxide synthase) and *pkm* (pyruvate kinase M) is inducible with hypoxia and hif1 targets showed a higher fold upregulation in female as compared to male.

**Fig. 2.**
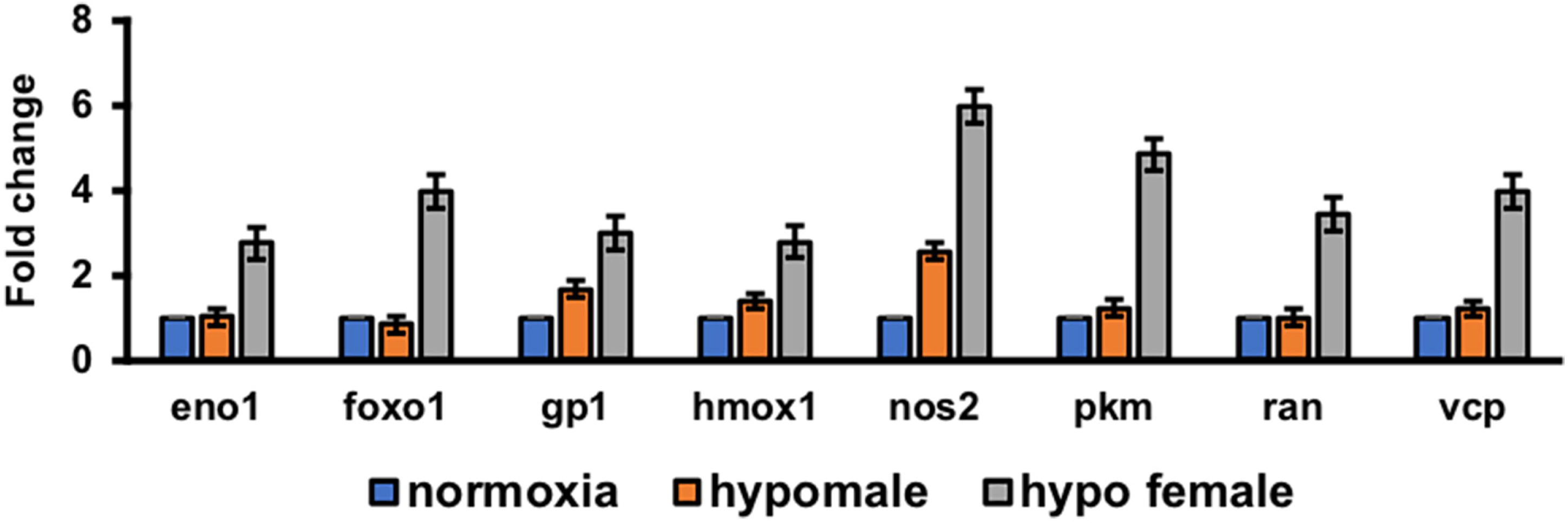
Validation of few regulatory target molecules. Graph showing mRNA expression of eno1, foxo1, gp1, hmox1, nos2, pkm, ran, and vcp. The data are expressed as the mean ± SEM, (n=6 pooled brains).

The core expression analysis of IPA led us to decipher many regulatory networks, disease function pathways, top upstream regulators and their predicted activation state, and also some biomarkers. Upon reviewing all the pathways involved, two individual pathways were found very interesting in male and female (Fig.3), which clearly showed gender-specific difference in the expression of a number of proteins in the pathway such as Rock1, Inppl1, Factin, Stat5ab, Ncor2, SRC-family, Got, Ints7 and Pdgfr, which were down regulated in male brain but upregulated in female brain. Though the pathways involved many molecules and networks, still were centred around the AKT signalling pathway, which regulates a wide range of cellular functions and is involved in the resistance response to hypoxia-ischemia through the activation of proteins associated with cell survival, proliferation and regulation of HIF-1α [23].

**Fig. 3.**
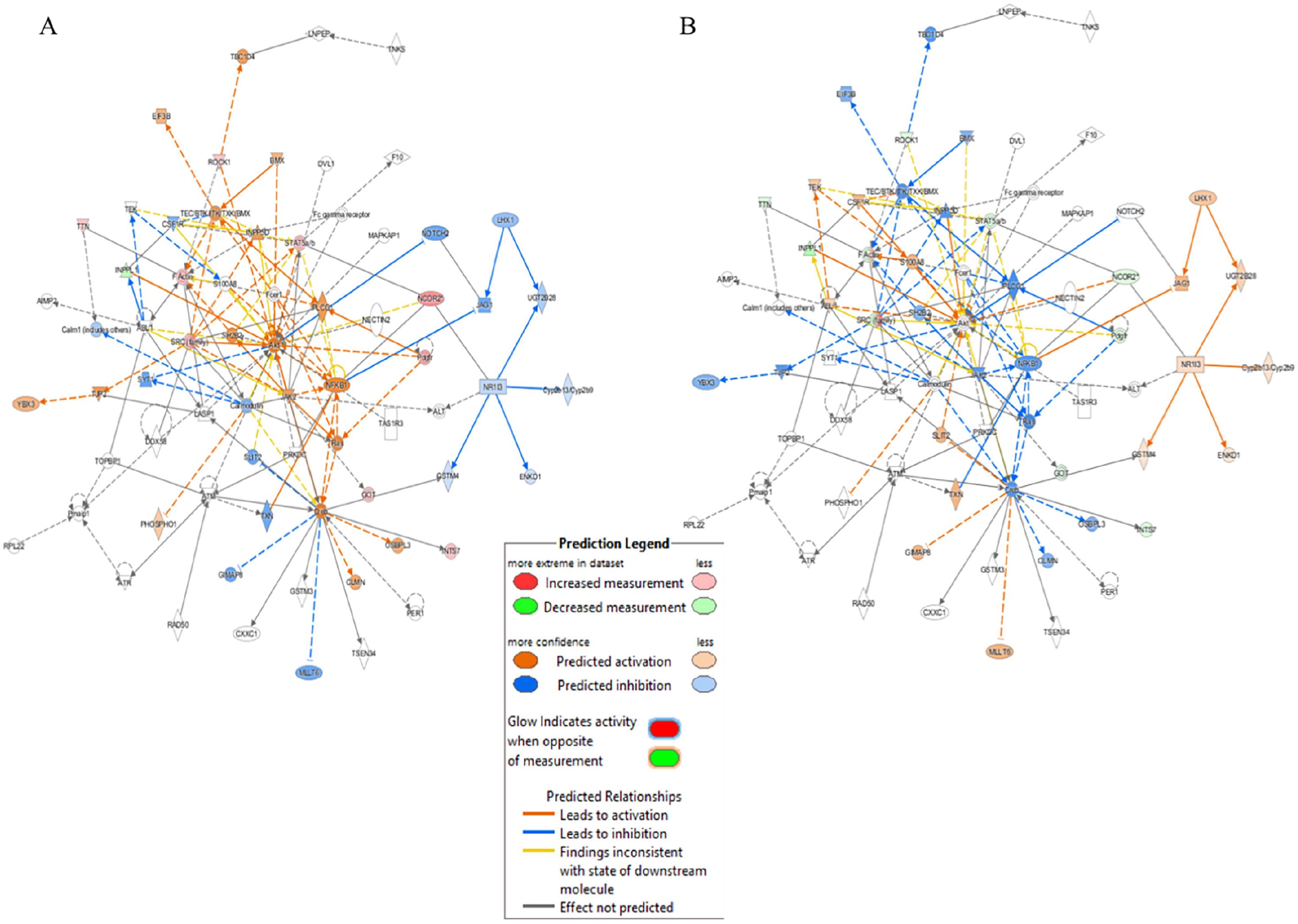
Pathway based on top networks generated by IPA. Predictive pathway for zebrafish male (A) and female (B) brain proteome induced by hypoxia

### Analysis of proteins showing uniform (either upregulated or downregulated) expression in both the sexes

Throughout the protein enrichment analysis by IPA sex-specific global proteome changes in the acute hypoxia zebrafish model was observed, which is in concurrence with our previous study [16]. But the question which remains unsolved is why the recovery in female quicker than in male. At 4 hr post-hypoxia when both the sexes survived coping up with the neural damage then there must be some common mechanism involved for recovery. So rather looking more into the differentially expressed markers we looked into common regulation of proteins. In Fig 1B we have shown the analysis of proteins resulted from iTRAQ where 554 proteins have common expression pattern in both the sex and among them 478 proteins were found regulated in one direction i.e. upregulated in both male and female brain in response to acute hypoxia. We hypothesized that as animals from both sex are in the recovery process therefore a common mechanism of regulation may help to elucidate the mechanism behind the late recovery of the male from neural damage induced by hypoxia-ischemia. While looking into the 478 upregulated proteins, among the top 5 upregulated proteins in male, we identified **histone-lysine N-methyltransferase H3** protein, an epigenetic regulator displaying ~3-fold upregulation in male brain and ~1.6-fold upregulation in female brain (Table 4.5). Post-translational modifications of histones are widely recognized as an important epigenetic mechanism in the organization of chromosomal domains and gene regulation. Methylation of lysine 4 and acetylation of lysine 9 of histone H3 have been associated with regions of active transcription, whereas methylation of H3K9 and H3K27 are generally associated with gene repression [24–28]. Recently hypoxia-induced histone modifications in neural gene regulation have been reported, and these were found on both hypoxia-activated and hypoxia-repressed genes [29]. H3K9 methylation is a critical epigenetic mark for gene repression and silencing. Hypoxia induces H3K9 methylation at different gene promoters, which is correlated with repression and silencing of those genes following hypoxia [30].

**Table 4.5.** Top 5 upregulated proteins retrieved from uniformly regulated (upregulated) in both male and female zebrafish brain induced by acute hypoxia.

| S. No. | Accession No. | Description | HM/NM | HF/NF |
| --- | --- | --- | --- | --- |
| 1 | 56693350 | Very long-chain acyl-CoA synthetase | 3.234 | 1.690 |
| 2 | **71834420** | **Histone-lysine N-methyltransferase, H3 lysine-9 specific 5** | **3.188** | **1.603** |
| 3 | 189531944 | PREDICTED: hypothetical protein LOC100148665 | 3.057 | 1.966 |
| 4 | 326666355 | PREDICTED: zinc finger protein 208-like | 2.984 | 2.606 |
| 5 | 326664965 | PREDICTED: protein FAM5C | 2.742 | 3.703 |

### Deciphering the role of H3K9me3 by co-immunoprecipitation and ChIP qPCR

Based on the previous literature [31], we hypothesized that H3K9 can be our prime target for deciphering late recovery in male as it was significantly upregulated in male brain following hypoxia and being a repressive epigenetic mark in nature its high level can repress and/or silence a number of critical neural genes. Considering the role of H3K9me3 in hypoxia [32] we immunoblotted for H3K9me3 using a specific antibody and performed a co-immunoprecipitation (CoIP) to identify the interacting proteins of H3K9me3 in hypoxic condition (Fig 4 A-C). We could validate the expression of H3K9me3 through immunoblotting with an upregulation of H3K9me3 in hypoxia male when compared to normoxia male (Fig.4A). For CoIP experiment nuclear extract was isolated from male zebrafish brain. The eluted proteins were then detected for immunoprecipitated and co-immunoprecipitated proteins by SDS-PAGE followed by western blotting. 5% of the initial lysates was used as the input (Fig 4 B). A mass spectrometric approach was used to identify the co-immunoprecipitated proteins obtained by the pull down of target antibody. The resultant peptides from MS/MS for four groups (normoxia IgG, normoxia and hypoxia H3k9me3 pull down) were analysed and after removing the background of IgG pooled proteins we could obtain 153 proteins identified in male normoxia H3K9me3 pull down and 72 proteins identified in male hypoxia H3K9me3 pull down. Surprisingly, there were no common proteins in normoxia and hypoxia H3K9me3 pull down, showing hypoxia stress may lead to alteration in interacting proteins. Further, we went through our iTRAQ data and tried to see whether these co-immunoprecipitated proteins were also found altered post-hypoxia in our high throughput proteomics data where almost all the proteins were found to overlap (Fig 4 C).

**Fig. 4.**
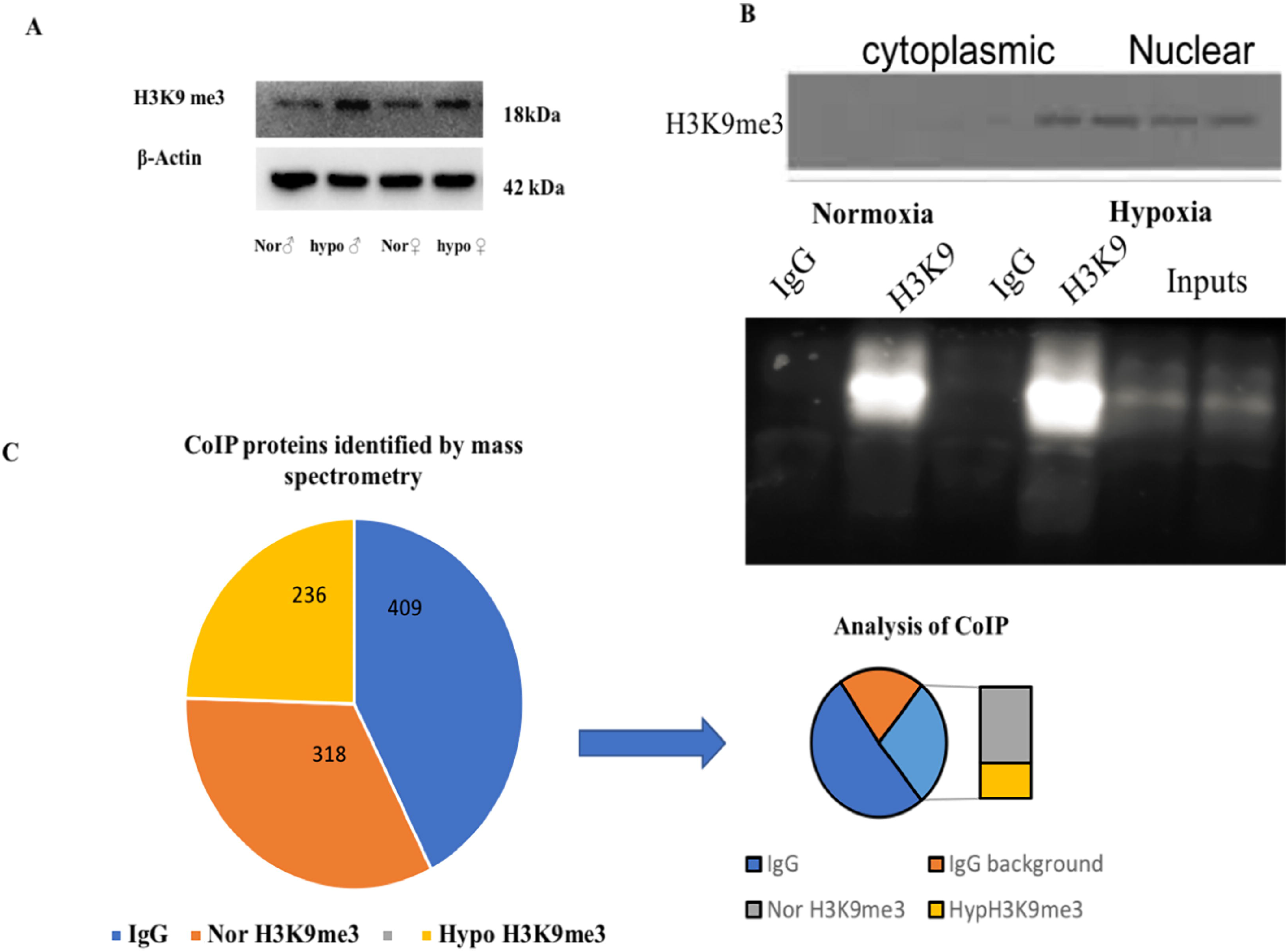
Deciphering the role of H3K9me3 by co-immunoprecipitation. Immunoblot of H3K9me3 showing upregulation in hypoxia (A), CoIP of H3K9me3 in zebrafish male brain nuclear protein (B), Schematic representation of CoIP analysis (C).

The co-interacted proteins of H3K9me3 could not answer the unresolved question why male is recovering late. Therefore, we thought of evaluating the transcriptional targets of H3K9me3 to get an answer to our question, as H3K9me3 is a repressive marker so its upregulation in male may repress any neurogenic marker needed for recovery from hypoxia induced neural damage. An earlier report on chromatin state of the developmentally regulated genes [31] led us to explore the striking upregulation of transcriptionally repressive epigenetic marker H3K9me3 at 4 hr post hypoxia in zebrafish male brain.

The ChIP-qPCR data showed the repression of early neurogenesis marker nestin, *klf4* and *sox2* in the zebrafish male brain 4 hr post-hypoxia. (Fig. 5) It is pertinent to mention here that the ChIP assay was not performed on female zebrafish brain as the male brain proteome showed a higher fold upregulation in H3K9me3. For further validating the data the mRNA expression levels for *nestin, klf4*, and *sox2* at two-time points of recovery i.e. at 4 hr and 12 hr post hypoxia, was assessed (Fig.6A-B). The qPCR analysis showed at 4 hr post-hypoxia the expression of early neurogenic markers showed mild activation in male and later at 12 hr post-hypoxia the expression was much higher. The protein level expression of Sox2 was evaluated at low concentration (25 microgram) of protein which showed in both male and female at 4 hr post-hypoxia but the expression was quite low in both the sex; in the male it was almost negligible, however in female a mild expression was observed, which at the later time point i.e. 12 hr post-hypoxia showed noticeable upregulation in both male and female brain.

**Fig. 5.**
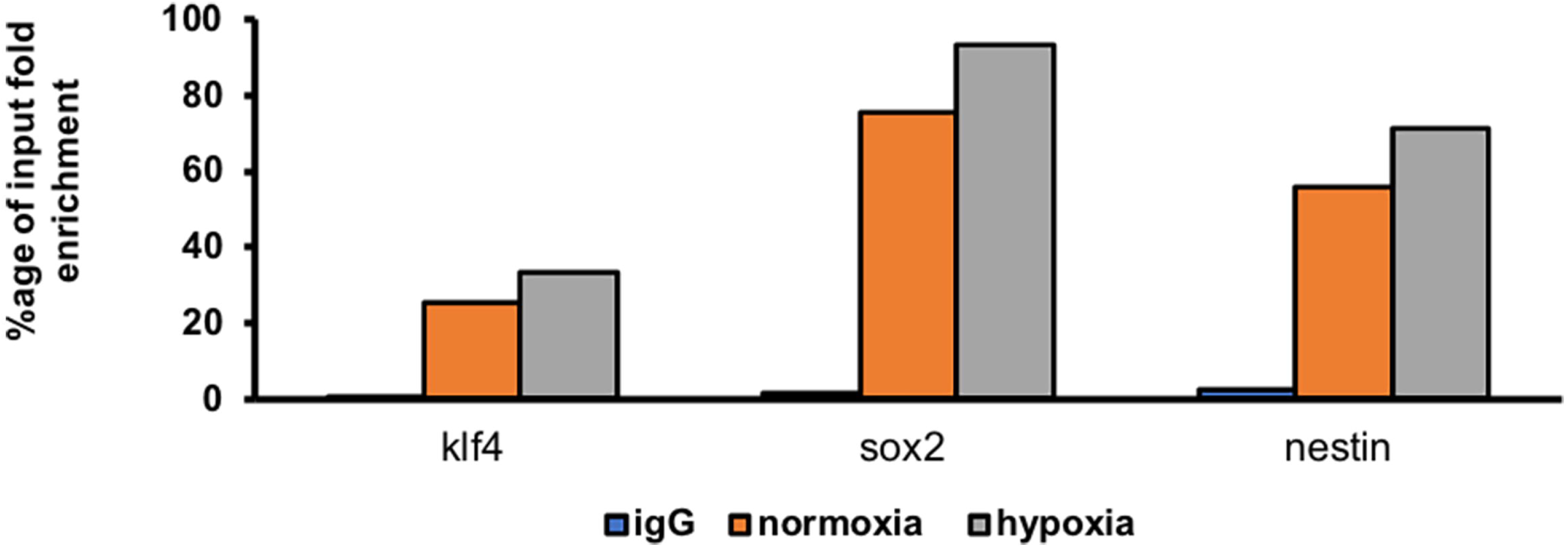
ChIP qPCR showing H3K9 occupancy on target gene promoter. ChIP qPCR showing the increase in H3K9me3 enrichment on the promoter region of klf4, sox2 and nestin in hypoxic male brain.

**Fig. 6.**
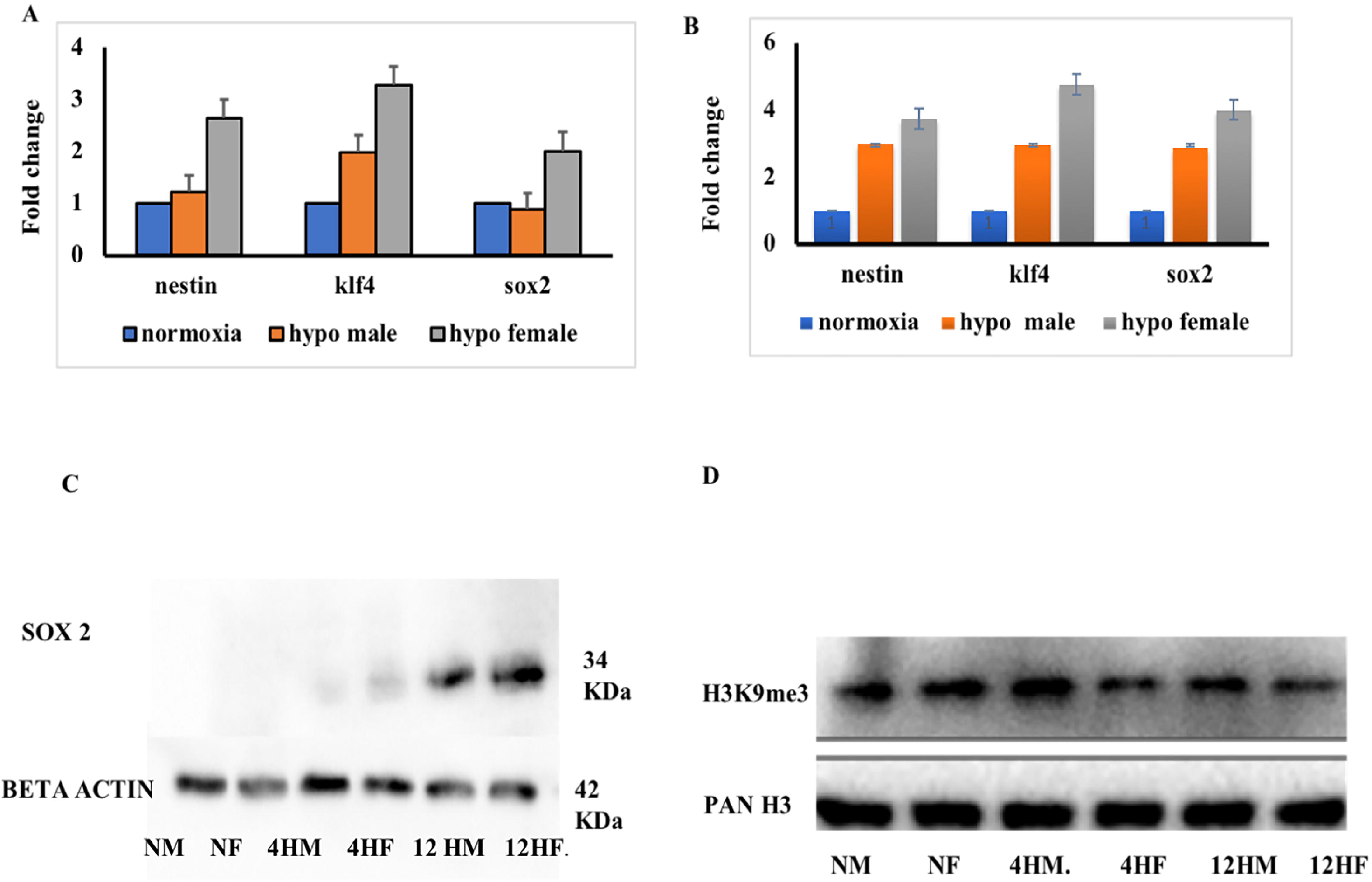
Validation ChIP analysis. mRNA expression at 4hr (A) and 10 hr (B) post hypoxia in both male and female (n=pooled 6 brain). Expression of sox2 protein at different time of recovery from hypoxia (C). Expression of H3K9me3 at different time of recovery from hypoxia (D).

To further validate if the sox2 expression is dependent on H3K9me3 level, the expression level of H3K9me3 was assessed and predictably it was found upregulated in male brain as shown in the previous experiment, compared to the female brain at 4 hr post-hypoxia. Later at 12 hr post-hypoxia, the level of H3K9me3 was much less in male than what it was at 4 hr post-hypoxia. This result suggested that with the activation of H3K9me3 the expression levels of early neurogenic markers are getting repressed. This could be the possible reason for late recovery in male as early neurogenic markers are not fully activated in response to hypoxia insult, in contrast to female brain.

Among Cerebral strokes, ischemic stroke is the most common type of stroke and major cause of death and/or disability worldwide, though there are continuous efforts to establish proper diagnosis and efficient therapy. The proteomics study complements both genomics and transcriptomics simultaneously provides information about the proteins that can implemented for main functional mediators of cells such as their post-translational modification and their interactions with biological molecules. However, post stroke is mostly related to protein function which can be direct target for therapeutic intervention. Therefore in the present study we performed a quantitative proteomics approach for hypoxia induced brain to identify favourable biomarkers involved in neuronal injury and recovery [10]. In our previous study clear sexspecific differences were observed in acute hypoxia-induced neural damage and recovery but to explore more about the mechanism of recovery in the present study we have focused at 4 hr post hypoxia timepoint predicting that could be possibly therapeutic window. To date, many high throughput studies on hypoxia [10, 33–39] gave sufficient information about the genes and proteins involved in hypoxia and related diseases but the roles of these hallmarked hypoxia markers are not well studied in a sex-specific context. Therefore, we have attempted to emphasize more on the sex-specific neural regulation post-hypoxia, which will provide a better insight into designing efficient therapeutics for patients who suffered acute hypoxic insults. The prevalence of hypoxic brain damage is increasing and prognostic factors for either poor or good outcome are lacking [40].

The advantage over traditional proteomics and iTRAQ based proteomics is in iTRAQ all the 4 groups can be simultaneously processed to reduce the error rate and post-translational modifications can also be quantified. The present study on whole zebrafish brain proteome upon global acute hypoxia sheds light on many differential roles of protein markers which can be further validated. Solute carrier (SLC) transporters are well known therapeutic targets [41] and in our study too we have observed a very high activation in recovery. We tried to elucidate the role of one of the histone based epigenetic regulatory mechanism (H3K9me3) that controls adult neurogenesis during the recovery phase post-hypoxia-ischemia. There is hardly any study on the epigenetic mechanisms in the zebrafish brain till date. Here, we identified hundreds of transcription factors involved in post-hypoxia recovery in a gender-specific manner, which can add to the development of better therapeutic strategy.

## CONCLUSION

To conclude we have studied the sex-specific difference in global proteome changes in zebrafish brain induced by acute hypoxia and during the recovery. We elucidated the unresolved question from our previous study [16] regarding the delayed recovery in male following hypoxic insult. With the striking upregulation of H3K9me3 in male at 4 hr post-hypoxia, the early neurogenic markers like nestin, klf4 and sox2 expression level got affected, which might be the reason for late recovery in male, compared to female. Acute hypoxia-induced sex-specific comparison of brain proteome led us to reveal many differentially expressed proteins including the novel ones, which can be further studied for the development of novel targets and a better therapeutic strategy.

## ACKNOWLEDGEMENT

This research was supported by the Council of Scientific and Industrial Research (CSIR), India network project (BSC0103-UNDO to SC and AK) and Department of Biotechnology, Government of India project (BT/PR14338/MED/30/495/2010 to SC). Authors thank the Director, CSIR-IICT for the support and the KIM Department for generating the official communication number (IICT/Pubs/2019/436).

